# Sinking the way: a dual role for CCR7 in collective leukocyte migration

**DOI:** 10.1101/2022.02.22.481445

**Authors:** Jonna Alanko, Mehmet Can Ucar, Nikola Canigova, Julian Stopp, Jan Schwarz, Jack Merrin, Edouard Hannezo, Michael Sixt

## Abstract

Immune responses crucially rely on the rapid and coordinated locomotion of leukocytes. While it is well established that single-cell migration is often guided by gradients of chemokines and other chemoattractants, it remains poorly understood how such gradients are generated, maintained and modulated. Combining experiment and theory on leukocyte chemotaxis guided by the G protein-coupled receptor (GPCR) CCR7, we demonstrate that in addition to its role as the sensory receptor that steers migration, CCR7 also acts as a generator and modulator of chemotactic gradients. Upon exposure to the CCR7 ligand CCL19, dendritic cells (DCs) effectively internalize the receptor and ligand as part of the canonical GPCR-desensitization response. We show that CCR7 internalization also acts as an effective sink for the chemoattractant, thereby dynamically shaping the spatio-temporal distribution of the chemokine. This mechanism drives complex collective migration patterns, enabling DCs to create or sharpen chemotactic gradients. We further show that these self-generated gradients can sustain the long-range guidance of DCs, adapt collective migration patterns to the size and geometry of the environment, as well as provide a guidance cue for other co-migrating cells. Such dual role of CCR7 as a GPCR that both senses and consumes its ligand can thus provide a novel mode of cellular self-organization.

## Main text

How cells navigate through tissues is one of the fundamental questions in biology, with great relevance in areas such as immunology, development, and cancer metastasis. Leukocytes, in particular, need to be recruited within short time frames because immunological threats arise at unpredictable sites in the body. Moreover, adaptive immunity is generated in lymphatic organs, which are usually remote from the primary site of infection, requiring long-range cellular trafficking^1,2^. A key concept derived from *in vitro* studies of cultured cells is chemotaxis, where cells migrate towards increasing concentrations of externally established, (supposedly) defined distributions of chemotactic cues. In a metazoan setting, the relationship between the chemoattractant distribution and cellular response is poorly understood. This is mainly because the spatio-temporal distribution of soluble chemotactic cues cannot be visualized *in vivo* and the cellular and molecular mechanisms that determine how gradients are generated and maintained in complex and dynamic tissues are largely unexplored^3–5^.

As a widely employed paradigm for leukocyte chemotaxis, we employed CCR7 guided migration of mature dendritic cells (DCs). Upon exposure to pathogenic threats, these cells rely on CCR7 to navigate from peripheral tissues towards the center of draining lymph nodes^6^. This process can be faithfully mimicked *in vitro:* when DCs are incorporated into 3D collagen gels overlaid with medium containing the soluble CCR7 ligand CCL19 (Fig.1a), the cells migrated directionally towards the chemokine diffusing from the overlaid compartment into the collagen gel (Fig.1b). Notably, cells showed a continuous increase in speed and persistence that lasted over hours and spanned millimeter distances (Fig.1b, Movie1). When we measured how the chemoattractant gradient evolves over time using 10kD FITC-dextran as a molecular proxy of comparable size, we found that the gradient gradually changed from a steep near-exponential distribution towards a flat, almost linear shape within 1-2 hours (Extended Data Fig.1a). While this might suggest that DCs migrate more efficiently in response to flat and linear gradients (Extended Data Fig.1b), this seemed unlikely based on previous findings that the chemotactic prowess of DCs is higher in exponential than in linear gradients^7^. Furthermore, immature DCs^8^ and other leukocytes like neutrophil granulocytes showed a very transient migratory response when exposed to chemoattractants in the same collagen gel setting (Extended Data Fig1c, Movie2). Such a response pattern was in line with the established model where cells can only effectively sense a gradient when the chemoattractant concentration is in the range of the Kd of the receptor and when the concentration difference of the ligand across the cell body (Δc/c) is sufficiently high to allow for spatial discrimination^9,10^. This, together with the long-recognized notion that the activity range of diffusion-based gradients of morphogens is very unlikely to reach the millimeter range^11^, led us to consider the option that DCs might re-shape the chemokine gradient as they migrate. Self-generated chemokine gradients have been demonstrated to drive the migration of cell clusters in the lateral line of the developing zebrafish. Here, cells at the leading edge express a sensory chemokine receptor that guides directed migration^12–15^. Cells at the trailing edge of the cluster express a separate, so-called decoy or scavenger receptor, which does not signal to G proteins but internalizes chemokines and thereby establishes a gradient. Furthermore, in some cancer cells and *Dictyostelium* amoeba, single cells carry cell surface enzymes that degrade guidance cues and have thereby been shown to organize collective locomotion^16–13^.

**Figure 1.**
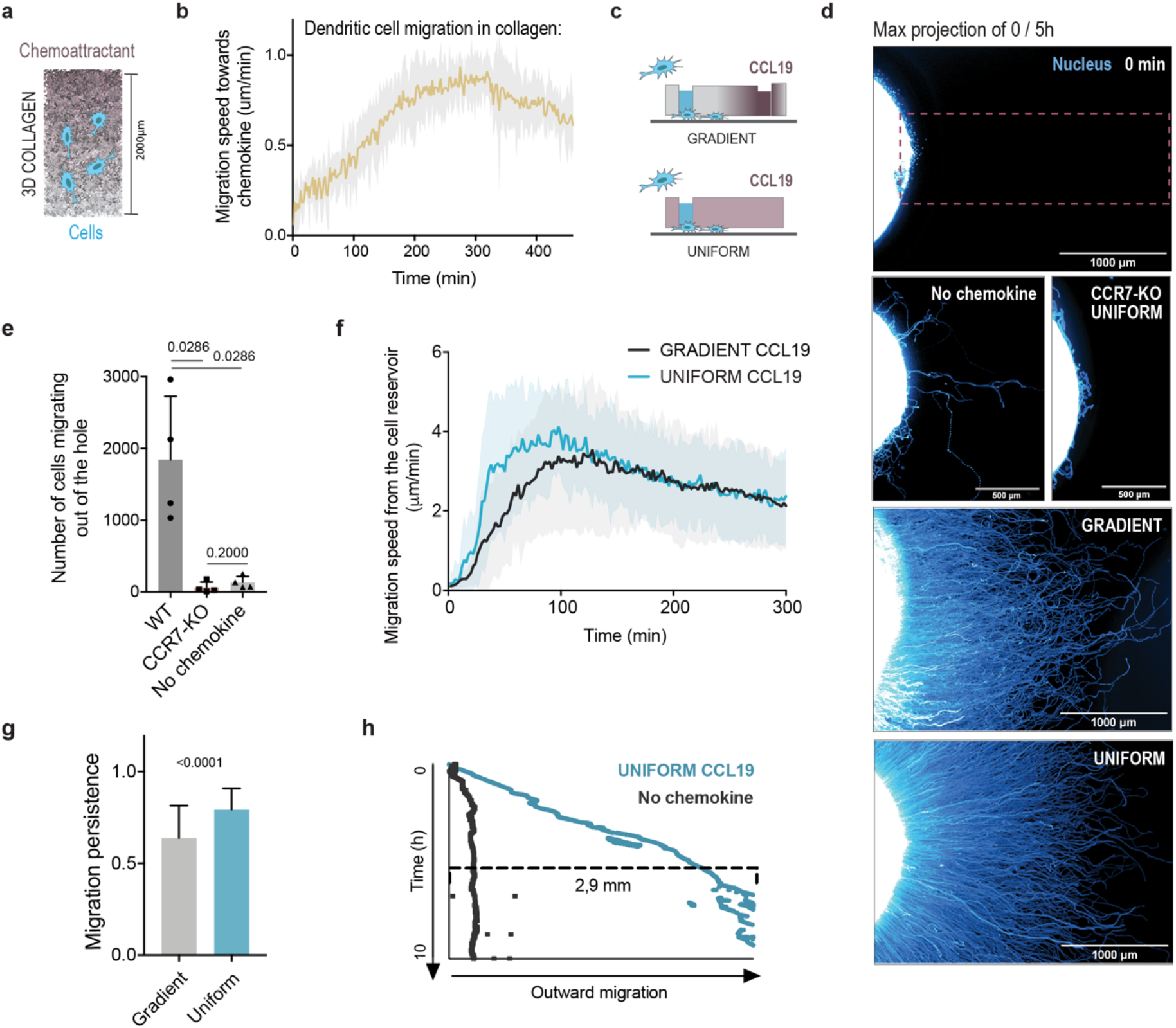
Dendritic cells migrate persistently and CCR7 dependently also in uniform CCL19 field. **a)** Schematic of 3D collagen assay with embedded cells and diffusible chemoattractant on top. **b)** Migration speed of mature dendritic cells (DCs) migrating in collagen towards CCL19 source in a population level for the first 7.5h (n=6 from 3 ind exp, mean+/-SD). **c)** Schematic of agarose assay, in which cells migrate from a reservoir to under agarose either towards a CCL19 reservoir or in uniform CCL19 field, where the same amount of CCL19 (18ng) is mixed with the agarose. **d)** Maximum projection of nuclear stain of WT or CCR7-KO DCs migrating out from the cell reservoir hole to under agarose for 0min or 5h with +/- CCL19 as gradient or uniformly. The dashed line box indicates the area used for quantifications in panels e,f,g. **e)** Number of cells migrating out of the cell reservoir under agarose with +/- uniform CCL19 (n=4 from 3 ind exp, mean+/-SD, Mann-Whitney test). **f)** Mean migration speed of DCs from the cell source during first 5h with a gradient or uniform CCL19 (n=5 from 3 ind exp, mean+/- SD). **g)** Migration persistence of single cells in the above under agarose experiments with uniform or gradient CCL19 (n(grad)=435 cells, n(unif)=1246 cells, mean ± SD, Mann-Withney test). **h)** Kymograph showing the progress of the leading edge of migrating DC population in +/- uniform CCL19 during the first 10h (representative of ≥3 ind exp).

To test if DCs shape gradients of their own chemoattractant, we used a system that allows modulating the spatial distribution of both cells and chemokine: we set up under agarose assays where DCs migrate in a confined space from a cell reservoir in response to different chemokine distributions (Fig.1c). In the absence of chemokine, only few cells showed persistent migration, while cells effectively and persistently moved towards a local source of CCL19 over a 5-hour imaging period (Fig.1d). Strikingly, similar numbers of DCs left the reservoir when CCL19 was offered uniformly. Both speed and directional persistence were even increased compared with the graded distribution, and cells in uniform concentration migrated persistently beyond the 3mm range (Fig.1f,g,h Movie3). CCR7 knockout cells barely left the reservoir, behaving similarly as cells without any chemokine (Fig.1d,e). These data strongly suggest that DCs create a gradient while they migrate, presumably by locally inactivating the chemokine.

While it was previously shown that DCs could proteolytically process the second CCR7 ligand CCL21, this was not the case for CCL19^19^. However, as part of the canonical GPCR desensitization pathway, CCR7 was shown to be rapidly endocytosed upon binding to CCL19 (Extended Data Fig.1d). For chemokine receptors, desensitization was previously discussed as a mechanism to terminate the migratory response whenever the local concentration exceeds a threshold, providing a signal to the cell to stop at its destination^2,20^. Alternatively, internalization followed by receptor recycling was suggested to clean receptors from ligands and thereby contribute to functional adaptation to high levels of ambient ligand^2,21^. We envisioned a third scenario, where internalization leads to consumption of the chemokine, a function that is well established for scavenger receptors but was not previously considered for canonical chemokine receptors.

When we visualized DCs upon exposure to labeled CCL19, we observed efficient uptake of CCL19, preferentially at the trailing edge of the cell, where CCL19 accumulated in small endosomal structures occasionally colocalizing with patches of actin (Fig.2a). Cells absorbed increasing amounts of CCL19 over time, and this was strictly CCR7 dependent, excluding the role of any unknown scavenger receptors in these cells (Fig.2b). CCL19 uptake could also be detected as chemokine depletion from the medium with a 5% decrease occurring during the first hour (Fig.2c). To test if this depletion was quantitatively sufficient to support a gradient-generating activity, we developed an under-agarose assay with two competing chemokine sources: one contained CCL19 only and the other CCL19 mixed with “sink” DCs. We then monitored the migration of small numbers of “sensor” DCs located in between both chemokine sources. When CCR7-KO DCs were used as sinks, sensor cells migrated equally towards both sources. When WT DCs were used as sinks, sensor cells migrated almost exclusively towards the cell-free source (Fig.2d), suggesting that the endocytic capacity of WT cells was sufficient to deplete CCL19 in functionally relevant quantities.

**Figure 2.**
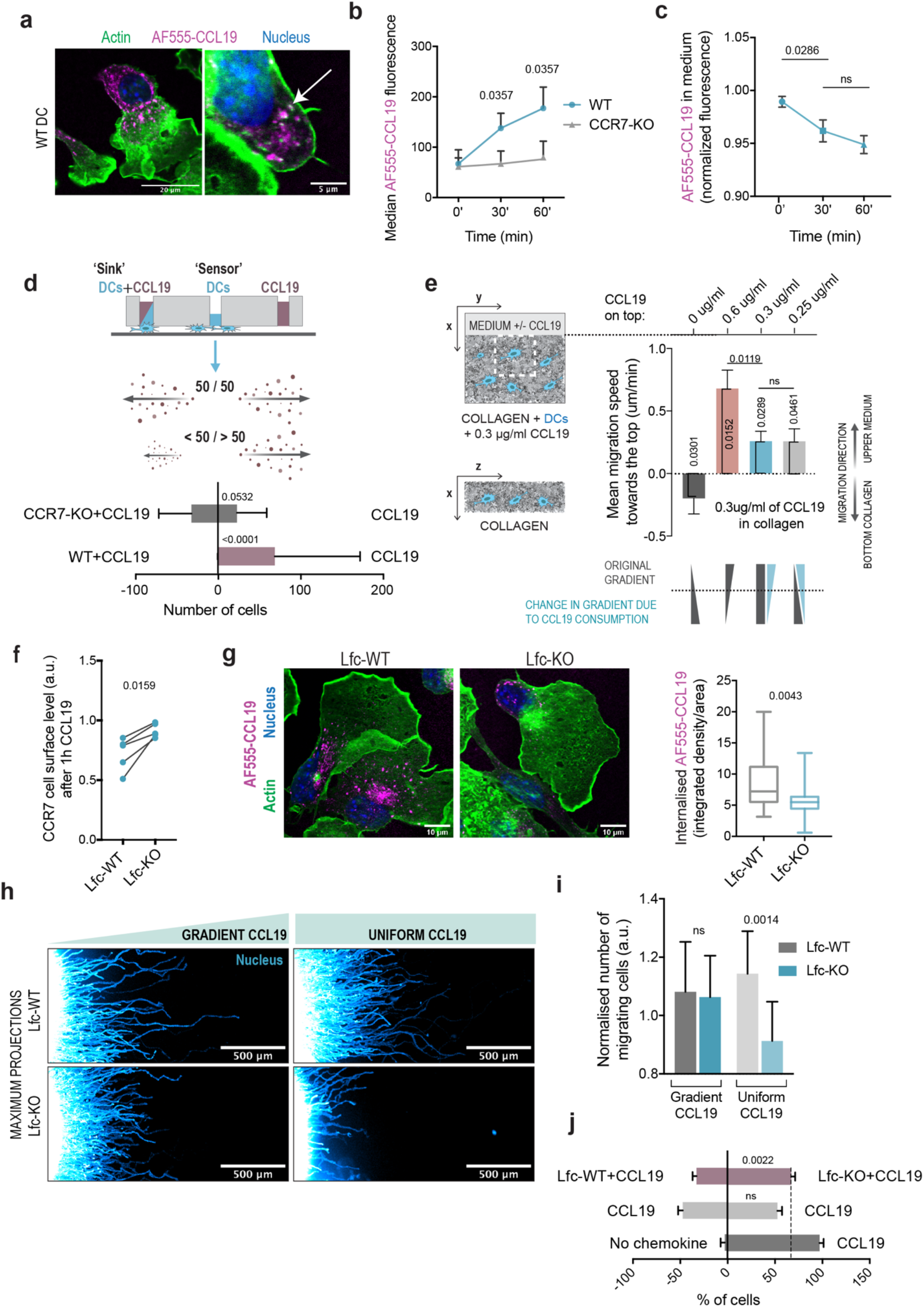
Dendritic cells create local CCL19 gradients by CCR7 mediated endocytosis. **a)** DCs incubated with AF555-labeled CCL19 under agarose for 2-3h, fixed and stained for nucleus (Dapi) and actin (phalloidin). The arrow points to an intracellular AF555-CCL19 vesicle-like structure positive for actin (representative images from 22 cells from 3 ind exp). **b)** Median AF555 fluorescence in WT and CCR7-KO DCs after incubation with AF555-CCL19 containing medium for 0, 30 or 60min (n=5,3 ind exp, mean+/-SD, Mann-Whitney test). **c)** Normalised AF555 fluorescence left in the AF555-CCL19 containing medium after incubation with WT DCs for 0, 30 or 60 min (n=4 ind exp, values normalised to the sum of all data points in an exp, mean+/-SD, Mann-Whitney test). **d)** Schematic of ‘Sink-Sensor’ under agarose setup to detect the ability of ‘Sink’ cells to deplete CCL19 from the medium by tracking the migration direction of the middle ‘Sensor’ cells. Below is a quantification of the number of ‘Sink’ cells migrating to left or right with WT or CCR7-KO DCs as the ‘Sink’ cells (n=15 tech rep from 3 ind exp. mean+/-SD, Mann-Whitney test). **e)** On the right is a schematic of 3D collagen setup, in which 0.3ug/ml of CCL19 is uniformly mixed with collagen and DCs, and medium containing different concentration of CCL19 is placed on top. The white dashed line box represents the detection field. On the left is quantification of the mean migration speed of DCs towards the upper compartment containing 0, 0.6, 0.3 or 0.25ug/ml of CCL19 during the first 8h (n=3 ind exp, mean+/-SD, values normalised to the sum of exp, one sample t-test (difference to 0 shown vertical, horizontal p-values from unpaired t-test). **f)** Antibody staining of CCR7 cell surface level in Lfc-WT and Lfc-KO BMDCs after 1h incubation with CCL19 (normalized to the corresponding 0min control, n=4 ind exp, Mann-Whitney test). **g)** Representative images of Lfc-WT and Lfc-KO BMDCs incubated with AF555-CCL19 for 2-3h before fixing and staining for nucleus and actin. On the right is quantification of the AF555-CCL19 signal inside the Lfc-WT and Lfc-KO BMDCs. Samples were imaged with LSM880 and the integrated density of AF555-CCL19 was measured from the middle slice/area of the cell (n=22, 26 cells from 3 ind exp, mean+/-SD, Mann-Whitney test). **h)** Maximum projections of stained nuclei in Lfc+/+ and Lfc-/- BMDCs migrating under agarose with gradient or uniform CCL19 for 10h (representative images of 3 ind exp). **i)** Number of migrating Lfc-WT and -KO BMDCs under agarose with gradient or uniform CCL19 (normalised to the sum of each experiment+0.8, mean+/-SD from 3 ind exp, paired t-test). **j)** Quantification of the % of ‘Sensor’ DCs migrating to left or right in above described ‘Sink and Sensor’ assay with +/- CCL19 on each side or Lfc-WT and Lfc-KO BMDCs as competing ‘Sinks’ (n=3,3,6 from 3 ind exp, mean+/- SD, Mann-Whitney test).

We next returned to the physiologically more relevant 3D collagen assays and tested if the sustained long-range migration of DCs towards CCL19 gradients might be caused by their gradient-amplifying activity. We embedded DCs in collagen gels containing uniform CCL19 and overlaid the gels with medium containing no DCs but varying concentrations of CCL19 (Fig.2e). As expected, DCs migrated towards the chemokine concentration, which was higher than the uniform level surrounding them. Importantly, cells overlaid with equal, or even lower, levels of CCL19 than the uniform CCL19 concentration persistently migrated to the overlaid chemokine source. This demonstrates that even when sparsely distributed in fibrillar 3D environments, DCs lower the environmental chemokine concentration sufficiently to collectively sustain their own chemotactic migration.

To challenge our hypothesis of internalization-mediated chemokine consumption, we aimed at compromising CCR7 endocytosis without altering its CCL19 binding capacity. Lfc is a Rho GEF recently shown to accumulate at the trailing edge of migrating DCs^22^, and the human homolog, GEH-F1 (Arhgef2), has been previously linked to vesicle trafficking^23,24^ and actin comet formation on endosomes (via RhoB)^25^. We found that Lfc-KO cells showed reduced CCL19-triggered CCR7 endocytosis (Fig.2f) and consequently decreased CCL19 uptake (Fig.2g). Directional persistence of Lfc-KO DCs was unimpaired when migrating in under agarose assays towards gradients of CCL19, indicating that the sensing function of CCR7 was intact (Fig.2h,i). In contrast, migration in uniform CCL19 concentration was substantially reduced. To confirm that this was due to insufficient gradient-generation activity, we used Lfc-KO and WT cells as “sink” cells in the aforementioned setup. Lfc-WT and -KO cells as competing sinks led the “sensor” DCs to migrate preferentially towards the Lfc-KO sink, confirming that CCL19 gradients are generated by endocytosis (Fig.2j).

To further challenge that endocytosis-mediated depletion of soluble chemoattractant could quantitatively explain the migration behavior detected in our agarose assays, we developed particle-based simulations for cells migrating in an initially uniform chemokine field. We modeled individual cells as soft polar particles performing a persistent random walk and able to degrade chemokine molecules in their proximity. Local self-generated chemokine gradients can furthermore be sensed by cells, causing them to reorient their polarity towards higher chemokine concentrations. Finally, the cells experience an effective repulsive force by other neighboring cells, representing steric interactions, see Supplementary Notes 1-3 and Supplementary Fig.3a for the implementation of model assumptions and estimation of parameters. This set of rules provided a minimal model to link individual cell motility and cell-chemokine interactions to the collective behavior of immune cell migration. For a wide range of parameter values, we systematically observed that cells migrated collectively in a propagating front away from their source (as observed before^26,27^), which robustly reproduced the experimentally obtained trajectories (Fig.3a, top panels). The simulations accurately captured the cell density and velocity profiles obtained from the experimental data (see Fig.3b), with velocities decaying over time due to flattening of the chemokine concentration, and densities propagating as a wave away from the cell source in comparable time scales as in the experiments. Simulations furthermore showed that cells in uniform attractant concentration are not able to migrate efficiently in case either attractant depletion or sensing is compromised (Fig.3a, bottom panels), in agreement with our CCR7-KO and Lfc-KO DC data.

**Figure 3.**
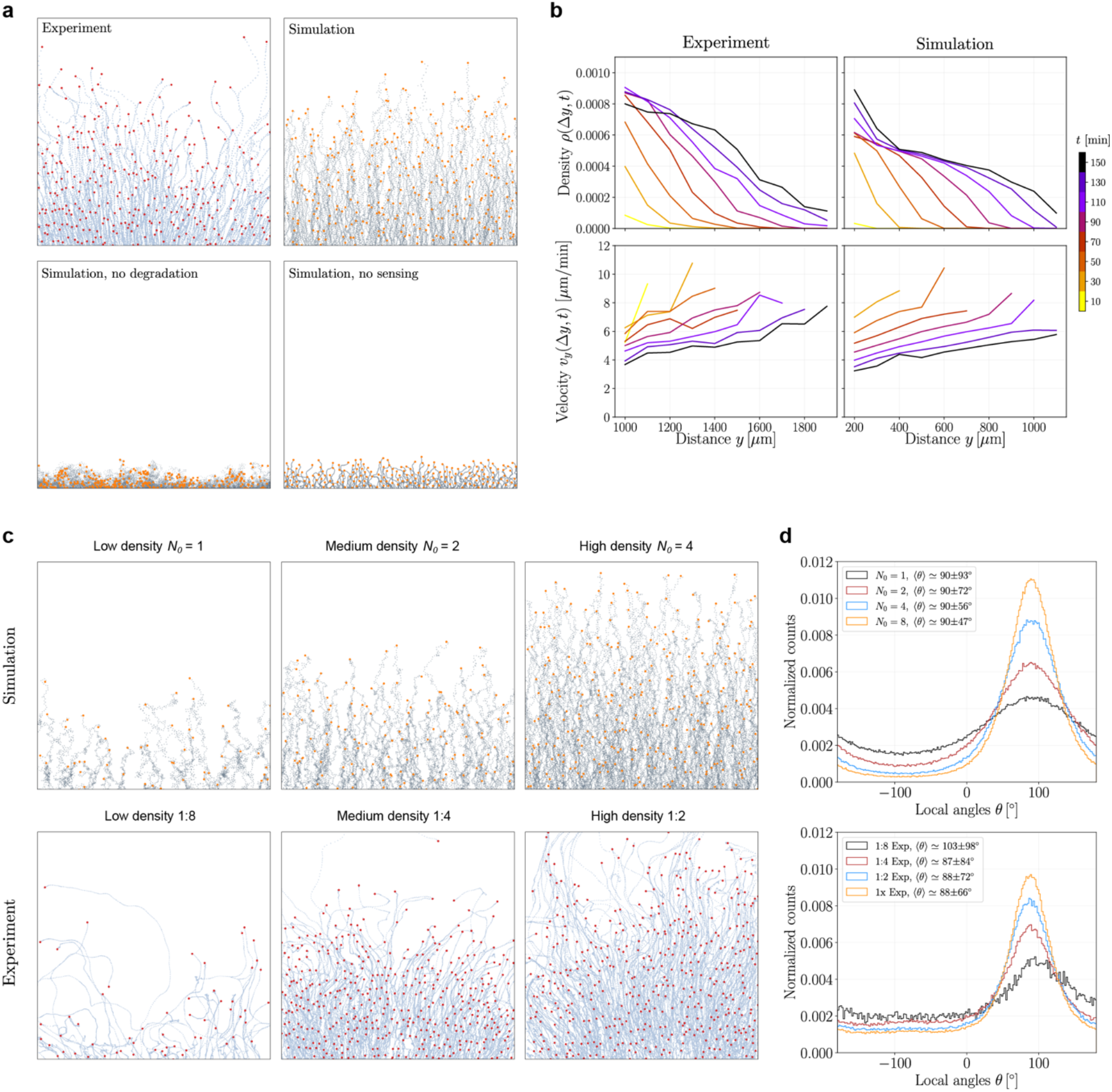
Particle based simulation shows directional bias being dependent on cell density and decreasing over time. **a)** Exemplary trajectories of DCs migrating in uniform CCL19 under agarose and in simulations (top panels) exhibit a qualitatively comparable bias away from the cell source. Bottom panels show simulations of cells that cannot deplete or additionally cannot sense chemokine molecules, leading to impaired migration (trajectories are representative of 4 ind exp and n=50 simulations). **b)** Spatio-temporal dynamics of cell densities and velocities as a function of distance from the cell source y, obtained from under agarose assays (left) and simulations (right), showing good agreement. Cell densities (top) propagate as a wave away from the source, with velocities (bottom) globally decaying over time (color-coded), while cells at the population front show consistently larger velocity. **c)** Exemplary trajectories of different densities of cells obtained from simulations (top) and under agarose assays (bottom). At a fixed propagation time, cells in higher density population migrate collectively over larger distances and exhibit a stronger directional bias. Densities are controlled by adjusting the cellular influx (number *N_0_* of cells added at fixed time intervals in simulations), and by different cell dilutions used in the experiments (denoted 1:2, 1:4 and 1:8). **d)** Directional bias of entire cell populations as obtained from the local angle (angle with respect to the horizontal axis) distributions of cellular trajectories in simulations (top) and experiments (bottom). Increasing the population density leads to narrower local angle distributions, with an average bias of *θ*-+90° and comparable fluctuations in simulations and experiments. 1x, 1:2, 1:4 and 1:8 are different cell dilutions.

Crucially, simulations also predicted that the directional bias depends strongly on the density of migrating cells, as denser populations of cells migrated over longer distances and more persistently (Fig.3c). We tested this experimentally with different amounts of DCs loaded into the cell reservoir and exposed to uniform CCL19, and found good qualitative agreement with the model prediction (Fig.3c, Extended Data Fig.1f). An intuitive mechanism for this feature is that single cells without followers create gradients in any given direction, leading to persistent yet unbiased overall migration, while cells in a collectively migrating group are effectively guided from the back, where the local chemokine concentration remains low due to internalization by follower cells^17,28^. As a result, cells at the population front face sharper gradients than those in the bulk, which lose their polarity over time as the local chemokine gradients become shallower. To probe the effect more quantitatively, we examined the angular distribution of migrating cells, with an angle close to +90° (resp. to −90°) indicating migration away from (resp. towards) the cell source. We found that the angular distributions became markedly sharper with increasing cell density (Fig.3c, right panels), with an approximately Gaussian shape having maxima centered around +90° and comparable standard deviations in experiments and simulations (Fig.3d). Finally, we found that the population front indeed exhibited increased collective polarity, as group polarization (a metric quantifying the collective alignment of active polar particles^29^) decreased over time as the population bulk advanced in the field of view (see Extended Data Fig.3d,e).

The ability of *Dictyostelium* amoeba to self-generate chemoattractive gradients was recently shown to allow them to solve geometric mazes^16^. To test the regulatory potential of CCR7 as a dual sensor and consumer of CCL19, we engineered a fork-like microfluidic device, where cells can choose between a closed channel and a channel leading to a big reservoir (Fig.4a). In the presence of uniform CCL19, WT DCs but not CCR7-KO DCs selectively entered and moved through the channel leading to the reservoir (Fig.4b, Movie4). This was in line with our theoretical simulations, showing that chemokine depletion from the closed channel favored entry into the reservoir-connected channel, where attractive gradients could build up (Fig.4c, Extended Data Fig.2d). The level of fluorescently labelled CCL19 was also detectably lower in the closed channel upon cell entry as well as behind a moving DC cluster, while the level inside the cells increased over time (Extended Data Fig.2a,b,c). These data demonstrate that CCL19 sinking endows DCs with the remarkable ability to assess environmental features at a distance.

**Figure 4.**
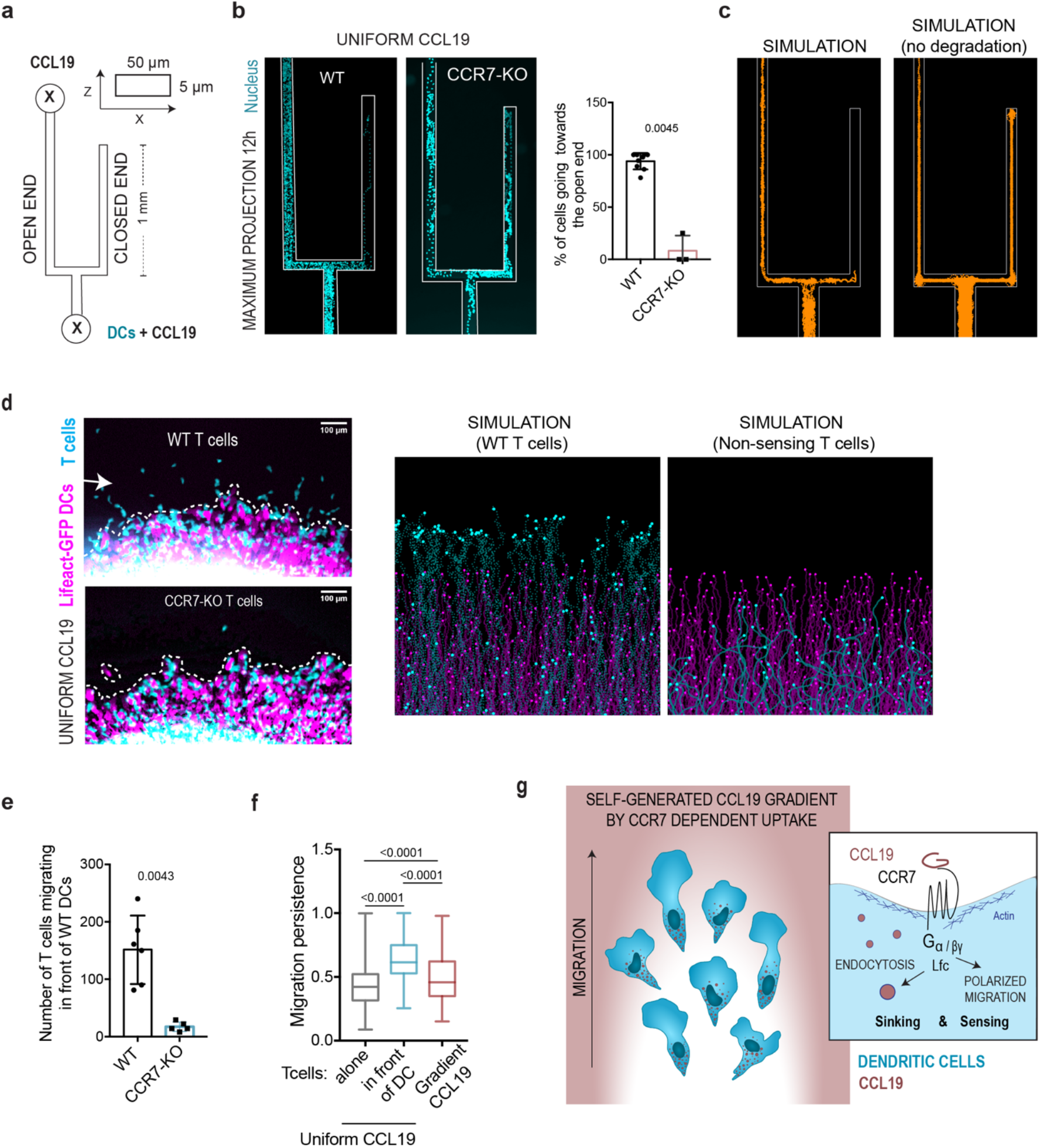
DC generated CCL19 gradients allow cells to avoid upcoming obstacles and guide other leukocytes. **a)** Schematic of the PDMS microfluidic device design with two inlets/reservoirs connected by 50*μ*m wide and 5*μ*m high channel. The whole device is filled with uniform CCL19, and cells are added in the lower inlet. **b)** WT and CCR7-KO DCs migrating for 12h in the device filled with uniform CCL19. Shown are maximum projections of nuclear stain, and quantification of the % of WT and CCR7-KO DCs migrating towards the open end of the device (mean+/-SD, n(WT)=9 from 3 ind exp, n(CCR7-KO)=3 from 2 ind exp). **c)** Corresponding cell migration trajectories obtained from simulations with (left) or without (right) attractant degradation. **d)** Snapshots of Lifeact-GFP expressing DCs (magenta) migrating together with TAMRA stained WT or CCR7-KO T cells (cyan) under agarose with uniform CCL19. White dashed line shows the front line of DCs and arrow examples of T cells migrating in front of DCs (representatives of 6,5 from 3 ind exp). **e)** Trajectories of DCs (magenta) and T cells (cyan) obtained from simulations exhibit an accumulation of T cells at a characteristic distance ahead of the DC front (left), while inhibiting the chemical sensing of T cells leads to an uncoupled migration profile of both cell types (right). **f)** Quantification of the number of T cells (WT or CCR7-KO) migrating in front of DCs (n=6,5 from 3 ind exp, mean+/- SD, Mann Whitney test). **g)** Migration persistence of WT T cells migrating under agarose with uniform CCL19 either alone or in front of DCs, or alone with gradient CCL19 (n>400cells from 3 ind exp, mean+/-SD, Mann Whitney test). **h)** Schematic of DC migration along self-generated CCL19 gradient. CCR7 has a dual role in chemokine sinking/depletion and sensing.

Finally, we asked whether CCR7-mediated self-generation of CCL19 gradients by DCs could affect the behavior of other cells. CCR7 guides pathogen-loaded DCs from inflamed peripheral tissues via lymphatic vessels into lymph nodes. T cells also use CCR7 to enter the lymph node, however, they arrive via the blood circulation, and they passage through lymph nodes not only upon inflammation but as part of their homeostatic trafficking program. Once in the same area, both cell types swarm around each other and establish short physical contacts to scan their surface receptors, eventually resulting in proliferation of antigen-specific T cells. However, whether and how DCs and T cells coordinate their local migratory response is not known. When we tested the response of either DCs or T cells to CCL19 under agarose, we found that T cells alone responded poorly to uniform CCL19 (Fig.4g). Upon mixing with DCs, T cells efficiently co-migrated with DCs in a CCR7 dependent manner (Fig.4d,f, Extended Data Fig.2e, Movie5). A wave of T cells preceding the migrating DCs up to roughly 150μm demonstrated that T cells could sense the CCL19 gradient that was generated by DCs but were unable or inefficient at generating their own gradient. This spatiotemporal structure of coupled wave propagation could be recapitulated in our numerical simulations by modeling T cells as a population unable to consume CCL19, but which could migrate with faster velocities than DCs in response to the same gradient (Fig.4e, Extended Data Fig.2f). Notably, the DC-generated CCL19 gradient was superior in guiding T cells compared to an externally applied CCL19 gradient (Fig.4g). These data suggest that DCs do not only have the capacity to orchestrate their collective migration to lymph nodes but can also guide the swarming behavior of T cells, which on their own have less or no capacity to modulate CCL19 distribution.

The dual role of CCR7 as sensor and sink (Fig.4h) does not only uncover a new level of functional chemokine regulation and cellular self-organisation with potentially wide-ranging physiological implications for immune cell communication, but it also raises questions regarding the role of receptor desensitisation as a way to functionally adapt a receptor to excessive ligand concentrations. While at the single-cell level, internalisation/recycling cleans the receptor from ligand and thereby allows for new binding events, we show that it also locally depletes the environment from ligands. This allows the cell collective to generate optimised concentration regime where the sensing of chemotactic gradients is effective because not all receptors are occupied with ligands. Evolutionarily, gradient-generating chemokine decoy receptors that lost G protein signalling but retained internalization capacity might be the specialized derivative of the dual function we describe here.

## Methods

### Mice

All C57BL/6J mice used in this study were bred and maintained at Institute of Science and Technology Austria’s animal facility in accordance with the institute’s ethics commission and Austrian law for animal experimentation. Permission for all experimental procedures was granted and approved by the Austrian Federal Ministry of Education, Science and Research (identification code: BMWF-66.018/0005-II/ 3b/2012).

### Chemokines

Purified recombinant mouse CCL19 (440-M3-025, R&D Systems) was used throughout the study. The Alexa Fluor 555 (AF555) labeled CCL19 was prepared with Alexa Fluor™ 555 Microscale Protein Labeling Kit (Invitrogen, A30007) according to manufacturers’ instructions. For the final elution, Zeba™ Spin Desalting Columns (7K MWCO, 89882, Thermo Scientific) were used, and the final protein concentration was measured with Nanodrop by using unlabeled CCL19 as a control.

### Cell culture

All cells were grown and maintained at +37°C in a humidified incubator with 5% CO_2_.

#### Dendritic cells (Hoxb8 and bone marrow-derived)

Dendritic cells were differentiated either from conditionally immortalized hematopoietic progenitor cells (described previously^30,31^) or from bone marrow of 8-11 weeks old female or male Lfc+/+ and Lfc-/- 129 mice (generous gift from Klaus-Dieter Fischer^22^). In brief, immortalized hematopoietic precursor cells were generated from wildtype, CCR7-/- or Lifeact-GFP C57BL/6J mouse bone marrow by retroviral delivery of an estrogen-regulated form of Hoxb8. Blasticidin selected cells were maintained in basic “R10” medium consisting of RPMI 1640 medium with 10% of heat-inactivated FBS, 100 U/ml penicillin, 100 μg/ml streptomycin and 50 μM β-mercaptoethanol (all from Gibco, Invitrogen) supplemented with 5% supernatant of an Flt3L producing cell line and 1 μM estrogen (Sigma, E2758). DC differentiation was induced by estrogen washout and culturing 2×10^6^ bone marrow cells or 4-5×10^5^ Hoxb8 cells in R10 supplemented with 10 or 20% of house generated granulocyte-macrophage colony stimulating factor (GM-CSF) hybridoma supernatant for 9 days (detailed protocol in^31^). After this, non-adherent cells were collected and DC maturation was induced by overnight incubation with 200ng/ml of lipopolysaccharide (LPS) from *E.coli* 0127:B8 (Sigma, L4516). Before every experiment, DCs were washed and allowed to recover at least for 1h in R10.

#### T cells

T cells were isolated from the spleen of C57BL/6J mice with EasySep Mouse T cell Isolation Kit (Stemcell, 19851) according to the manufacturer’s instructions. Isolated T cells were then activated by plating the cells on cell-culture wells coated for overnight with 1 μg/ml of antibodies against CD3e and CD28 (16-0031-82 and 16-0281-82, Invitrogen) for 2 days in R10 medium supplemented with IL-2 (10ng/ml, 402-ML-020, R&D Systems) for 2 days. After this, activated T cells were collected and expanded in IL-2 containing R10.

#### Neutrophil-like cells

Neutrophil-like HL-60 (a gift from Alba Diz-Muñoz) and PLB-985 cells (from DSMZ, ACC 139) were maintained in RPMI 1640 supplemented with 10% FBS, 20 mM HEPES and 1% glutaMAX (all Gibco, Thermo Fisher Scientific). For differentiation, cells were kept for 6 days in medium containing 1.25% DMSO (cell-culture grade; Sigma-Aldrich), and the differentiation status was validated using flow cytometry (CD11b, Miltenyi Biotec). Before experimentation, differentiated cells were washed three times to remove DMSO.

### Migration assays

#### In collagen

3×10^5^ DCs or 1.5×10^5^ differentiated neutrophil-like cells were mixed with collagen I solution containing 1.7mg/ml of purified bovine collagen I (5005, 5005, advanced Biomatrix), 1x of minimum essential medium (21430020, Gibco) and 0.3% of sodium bicarbonate (NaHCO_3_ s8761, Sigma) in R10 medium (total volume 300μl). The collagen-cell mixture was then casted in custom-made chamber and collagen was allowed to polymerize for 45min at 37°C before overlaying the gel with either 0.6ug/ml of recombinant mouse CCL19 (for DCs, 440-M3-025, R&D Systems), 25nM fMLP (for neutrophil-like cells, F3506-5MG, Sigma) or with 10kD FITC-dextran. In case of FITC-dextran diffusion, the assay was done without collagen embedded cells. The chamber was sealed with paraffin and cell migration was recorded immediately with time-lapse video microscopy for 4-8h at 37°C, 5% CO_2_ with 60s intervals. The average migration speed of the entire cell population towards the chemokine source was determined with a custom-made ImageJ plugin (by Robert Hauschild, Institute of Science and Technology Austria). In brief, images were background corrected and particle filtering was used to discard objects other than cells. For each frame, the lateral displacement optimizing its overlap with the previous frame was determined and divided by the time interval between the frames, yielding the y-directed migration speed of the cell population over time. The respective script can be shared upon request. In case of uniform CCL19, 0.3μg/ml of CCL19 was mixed with collagen and DCs before polymerization, after which the gel was overlaid with different concentrations of CCL19.

#### Under agarose

400μl of agarose mix containing 1% of UltraPure™ Agarose (16500100, Invitrogen) in phenol-free R10 medium supplemented with 1x Hanks’ buffered salt solution (pH 7.3) and 10% FBS was poured on custom-made chambers with diameter of 17mm and allowed to polymerize for 35min. In case of gradient chemokine, two holes (diameter 2mm) were punched 1cm apart into the agarose for 0.5-1 μl of pelleted cells and for 17.5ng of CCL19 in phenol-free R10 for a chemokine source. For uniform chemokine concentration, 17.5ng of CCL19 was mixed with the agarose mix before polymerization and only one hole was punched into the agarose. Cells were stained with NucBlue™ (R37605, Invitrogen) for 15min at 37°C and washed once with R10 before injecting into the agarose hole. Cell migration was recorded with inverted Nikon Ti widefield fluorescent microscope at 37°C with 5% CO_2_ for 6-12h with 1min interval and 4x or 10x objective.

*For the ‘Sink and Sensor’ setup,* three 1.5mm holes 5mm apart were punched into the agarose. 2.5μl of pelleted DCs were mixed with 10ng of CCL19 in colorless R10 with total volume of 4μl, and injected into one of the edge holes. For chemokine-only controls, 10ng/4μl of CCL19 in colorless R10 was used. 15 000 NucBlue stained DCs in 4μl of colorless R10 were injected into the middle hole as the ‘Sensor’ cells and the migration of these cells was recorded with fluorescent microscope. In the case of Lfc+/+ and Lfc-/- BMDCs, cells were incubated with 10ng of CCL19 at 37°C for 3h before injection into the agarose hole.

#### T cell migration together with DCs

T cells were stained with 10μM 5-carboxytetramethylrhodamine, succinimidyl ester (TAMRA, Molecular Probes, Life Technologies) for 10min RT, washed with R10 and allowed to recover for 30min before use. Both Lifeact-GFP DCs and T cells were also stained with NucBlue. Equal number of both cell types were mixed together and pipetted into a 2mm hole in agarose. The gel was prepared otherwise as above, but with 0.75% of UltraPure™ Agarose (16500100, Invitrogen). Cells were imaged with inverted Nikon Ti widefield fluorescent microscope at 37°C with 5% CO_2_ for 3h with 1min interval and 10x objective. To quantify T cell migration in front of DCs, the Lifeact-GFP-expressing DCs were selected by filtering (variance 30) and thresholding, and this selection was used to clear out T cells interacting with DCs from the T cell/555 channel, thereby leading to a movie showing only T cells migrating front of DCs, which was then used for tracking with Trackmate.

### Image analysis & statistics

Fiji^32^ was used for all image processing and analysis in this work, and all graphs and statistical analyses were done with GraphPad Prism. The number of independent biological replicates and experiments used for the graphs as well as used statistical test are indicated in the figure legends. For most results mean+/- standard deviation (SD) was used with Mann-Whitney test.

#### Quantification of cell migration

Cell migration was quantified by using Trackmate plugin for automated cell tracking^33^ with carefully adjusted parameters and filtering to exclude unmoving dead cells and bad tracks. The same parameters were used consistently in different conditions to obtain comparable results. The NucBlue channel (405) was used for the tracking. The cell reservoir was cleared from the time-lapse images by appropriate filtering and thresholding before tracking.

The kymograph showing the progress of the leading edge of migrating DC population (in Fig.1h) was created by using variance filtering, thresholding and finding the edge followed by reslicing to obtain a kymograph.

#### Quantification of internalized AF555-CCL19 from immunofluorescence images

The integrated density of AF555-CCL19 inside the cells (Fig.2g) was measured from the middle slice by manually drawing a selection based on the actin staining. For the quantification, the measured integrated density was normalized to the corresponding cell area.

#### Quantification of DC length

The mean length of a migrating DC under agarose (used for the model) was measured manually from brightfield images.

### Immunofluorescence

DCs were allowed to migrate under agarose with uniform AF555-CCL19 for 2-3h before fixing with prewarmed (37°C) 4% paraformaldehyde (PFA, e15714, Science services) for 15min at RT. The agarose layer was removed, PFA was washed away 3x with PBS, and cells were stained with Dapi and Alexa Fluor™ 488 phalloidin (A12379, Invitrogen) for 1h at RT, followed by 3x PBS wash and mounting with ProLong™ Gold Antifade Mountant (P36930, Invitrogen). Samples were imaged with Zeiss LSM880 upright or inverted microscopes with Airy Scanner with Plan-Apochromat 40x / NA 1.2 Water or 40x / NA 1.4 Oil objective.

#### Uptake of AF555-CCL19

300 000 DCs were incubated with 100ng of AF555-CCL19 in 100 l of colorless R10 for 0, 30 or 60min at 37°C. Cells were then transferred on ice and the AF555 fluorescence in cells was measured with FACS CANTO II flow cytometer (BD Biosciences). Cells without any AF555-CCL19 were used as a negative control and this value was subtracted from the rest. To measure the level of AF555-CCL19 in the medium, 600 000 DCs were incubated with 11ng of AF555-CCL19 in 100*μ*l of colorless R10 for 0, 30 or 60min at 37°C, after which the samples were centrifuged (300xg, 5min) and the AF555 fluorescence of the supernatants was measured with Biotek SynergyH1 plate reader. Colorless R10 medium was used as a negative control. The measured fluorescence values were normalized to the sum of the corresponding experiment and multiplied by 3 to reduce the variation between independent experiments.

#### Flow cytometry to measure CCR7 cell surface level

300 000-400 000 cells were incubated with different concentration of CCL19 in 100ul of R10 for different time points at 37°C in 96 well, after which cells were fixed by adding equal volume of 4% PFA on ice for 15min. Cells were washed 2x with PBS and stored in FACS buffer (2mM EDTA, 1% BSA in PBS) at 4°C or washed with FACS buffer and immediately stained. Cells were incubated first with 5*μ*g/ml of α-CD16/CD32 (14-0161-85, eBioscience) in FACS buffer for 10min at RT to block FC receptors. Cell surface CCR7 was stained by incubating the cells in 1:300 of mCCR7-PE antibody (4B12, 12-1971-82, eBioscience) in FACS buffer containing the FC block for 1h at RT, after which cells were washed 3x with FACS buffer before measuring the median PE signal with FACS CANTO II flow cytometer (BD Biosciences). For the quantification in Fig.2f, the measured median fluorescence values were normalized to the corresponding 0min CCL19 control.

### Simulation

Details of the particle-based model, simulation settings for comparison with the under agarose assays, microfluidic device experiments, and for heterogenous cell populations, as well as parameter values used in simulations can be found as a separate Supplementary text file.

### Migration in micro-fabricated devices

The microfluidic design was drawn in CorelDraw 2019, converted to Gerber format with LinkCad, and printed on a 5” chrome photomask (JD Photo Data). The 5-micron high master wafer was made by UV photolithography using SU-8 TF 6005 photoresist (Kayaku Advanced Materials) and characterized with a profilometer. The master wafer was then vapor treated with 1H,1H,2H,2H-perfluorooctyl-thichlorosilane for one hour. Polydimethylsiloxane (PDMS) devices were made with Sylgard 184 Elastomer Kit (Dow Corning) using a ratio of 10:1. Mixed PMDS was poured on the wafer, degassed in a vacuum chamber, and baked overnight at 80°C. The hardened PDMS layer was then peeled off, cut into single devices, and through holes of 2mm diameter were punched out. Finally, the devices were bonded onto EtOH-washed coverslips by oxygen plasma followed by sealing at 85°C for 15min to make the bond permanent. The finished PDMS device with 50×5*μ*m channels was filled and preincubated with R10 for overnight at 37°C, after which the device reservoirs were emptied, and the whole device was filled with either 2.5*μ*g/ml of CCL19 or 11*μ*g/ml of AF555-CCL19 in colorless R10. The chemokine was allowed to equilibrate for a minimum of 3h before the addition of 0.5-1ul of pelleted NucBlue stained DCs to the cell reservoir. Cells were imaged every 1min with an inverted Nikon Ti widefield fluorescent microscope at 37°C with 5% CO_2_ for 12h and 4x objective. The number of cells migrating towards the open-end and closed-end was manually counted.

## Supporting information

Supplementary Theory Note

## Acknowledgements

The authors thank the Scientific Service Units (Life Sciences, Bioimaging, Nanofabrication, Preclinical and Miba Machine Shop) of the Institute of Science and Technology Austria for excellent support. This work was supported by grants from the European Research Council under the European Union’s Horizon 2020 research to M.S. (grant agreement No. 724373) and to E.H. (grant agreement No. 851288), and a grant by the Austrian Science Fund (DK Nanocell W1250-B20) to M.S. J.A. received funding from Jenny and Antti Wihuri Foundation. M.C.U. was supported by the European Union’s Horizon 2020 research and innovation programme under the Marie Skłodowska-Curie grant agreement No 754411.

## Author Contributions

J.A. and M.S. conceived the experiments and wrote the manuscript together with E.H. and M.C.U. N.C. reviewed and corrected the manuscript. N.C. performed and analyzed the experiment with sink and sensor DCs, J.St. performed and analyzed the collagen migration assays with neutrophils, and J.Sc. performed the FITC dextran diffusion experiment. All other experiments were performed and analyzed by J.A. The wafers for PDMS devices were designed and prepared by J.M.

M.C.U. and E.H. developed the particle-based model. M.C.U performed the simulations and analyzed the simulation and experimental data.

## Data availability

Authors can confirm that all relevant data are included in the article and/or its supplementary information files. Original videos will be made available at a reasonable request due to the large file size. Raw trajectory data used to compare the simulations and experiments are available from the authors upon request.

## Code availability

Custom-made scripts to reproduce the particle-based simulations and analyze the data are available from the authors on request.

## Supplementary Information

is available for this paper.

---

**Extended Data Figure 1.**
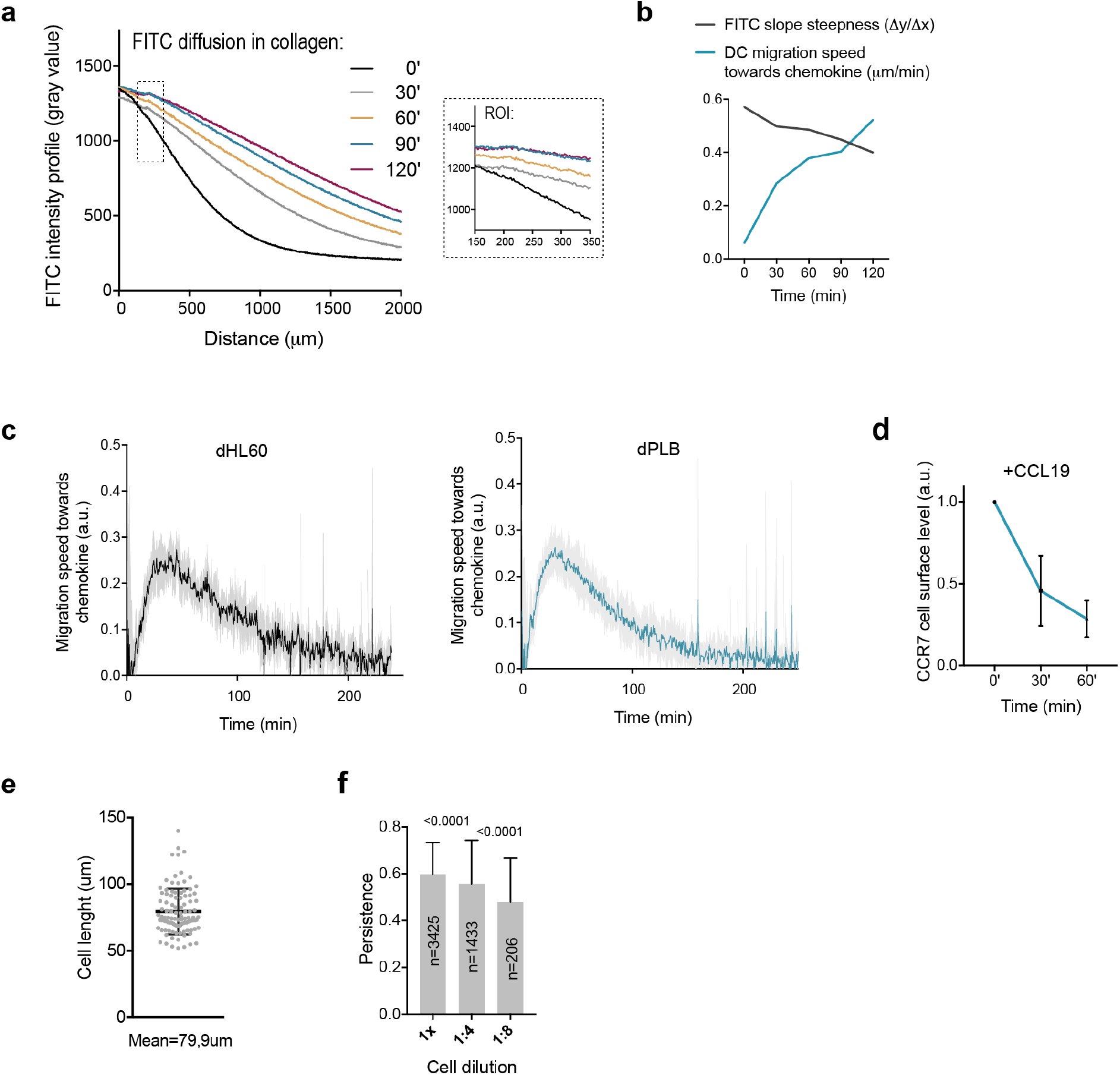
**a)** Diffusion of 10kD FITC-dextran into 3D collagen over time. Shown are intensity profiles at different time points (0, 30, 60, 90 and 120min) in a 2000um region (similar to the cell assays) and a close up (ROI) from a shorter distance (representative of 2 ind exp). **b)** A graph showing DC migration speed in 3D collagen (in Figure 1b) over the first 120min (um/min) and the slope steepness of FITC intensity profile. **c)** Migration speed of neutrophil-like dHL60 and dPLB cells towards 25nM fMLP in 3D collagen setup over time (mean+/-SD, 5 ind exp). **d)** Antibody staining of CCR7 cell surface level in DCs incubated with 250ng/ml of CCL19 for 0, 30 or 60min (n=2-3 from 3 ind exp, mean+/-SD). **e)** Quantification of the average length of DC migrating under agarose with uniform CCL19 (n=381 cells from 4 exp, 3 ind exp, mean+/-SD). **f)** Persistence of DCs (1x, 1:4 and 1:8 dilutions) migrating under agarose with uniform CCL19 in different dilutions (1x, 1:4 and 1:8) (from 2mm hole) (n=3425, 1433, 206 cells from 3 ind exp, mean+/-SD, Mann Whitney test).

**Extended Data Figure 2.**
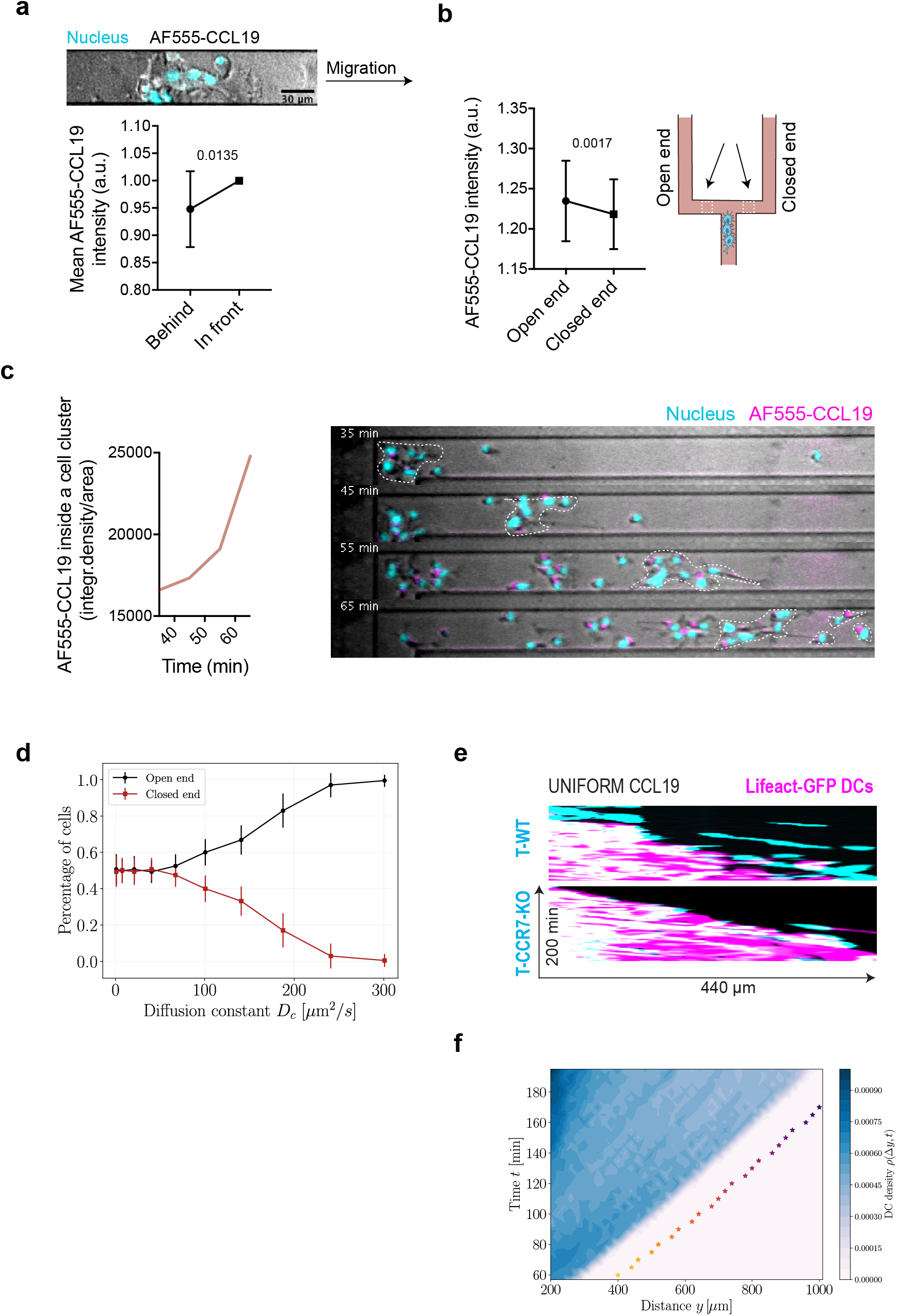
**a)** Example of a DC cluster migrating in microfluidic channel with uniform CCL19 (nuclei in cyan, AF555-CCL19 in white) and quantification of normalized mean intensity of AF555-CCL19 in microfluidic channel in front and behind of a migrating cell cluster (n=10 clusters from 4 samples, 2 ind exp, mean+/-SD, Mann Whitney test). **b)** AF555-CCL19 intensity (normalized to the corresponding background value) in the microfluidic channel on the side of open-end and closed-end channels, and a schematic illustrating the sites of measurements (while dashed line boxes) (n=8 measurements from 4 samples, 2 ind exp, mean+/-SD, paired t-test). **c)** Quantification of the increasing AF555-CCL19 intensity (integrated density/area) inside a moving DC cluster over time in a microfluidic channel filled with uniform AF555-CCL19 and example snapshots. The white dashed line indicates the followed cell cluster used for measurements (nuclei in cyan, AF555-CCL19 in magenta, representative of 4 samples from 2 ind exp). **d)** Predictions from particle-based simulations on the percentage of cells entering the open-(black) vs. the closed-end (red) channels as a function of the diffusion coefficient of the chemokine. The probability of cells entering the open-end channel increases monotonically with increasing chemokine diffusibility, where values of 100-200 um^2^/s correspond to the experimentally relevant estimates. **e)** Kymographs of experiments shown in Figure 4d (the first 200min). **f)** Kymographs of DCs and T cells as obtained from the simulations. Color bar denote the DC densities, and the star symbols represent the concentration maxima of T cells at different time points (color-coded). Despite their larger chemotactic sensitivity, T cells preserve a characteristic length scale of ~150um at all times.

**Extended Data Figure 3.**
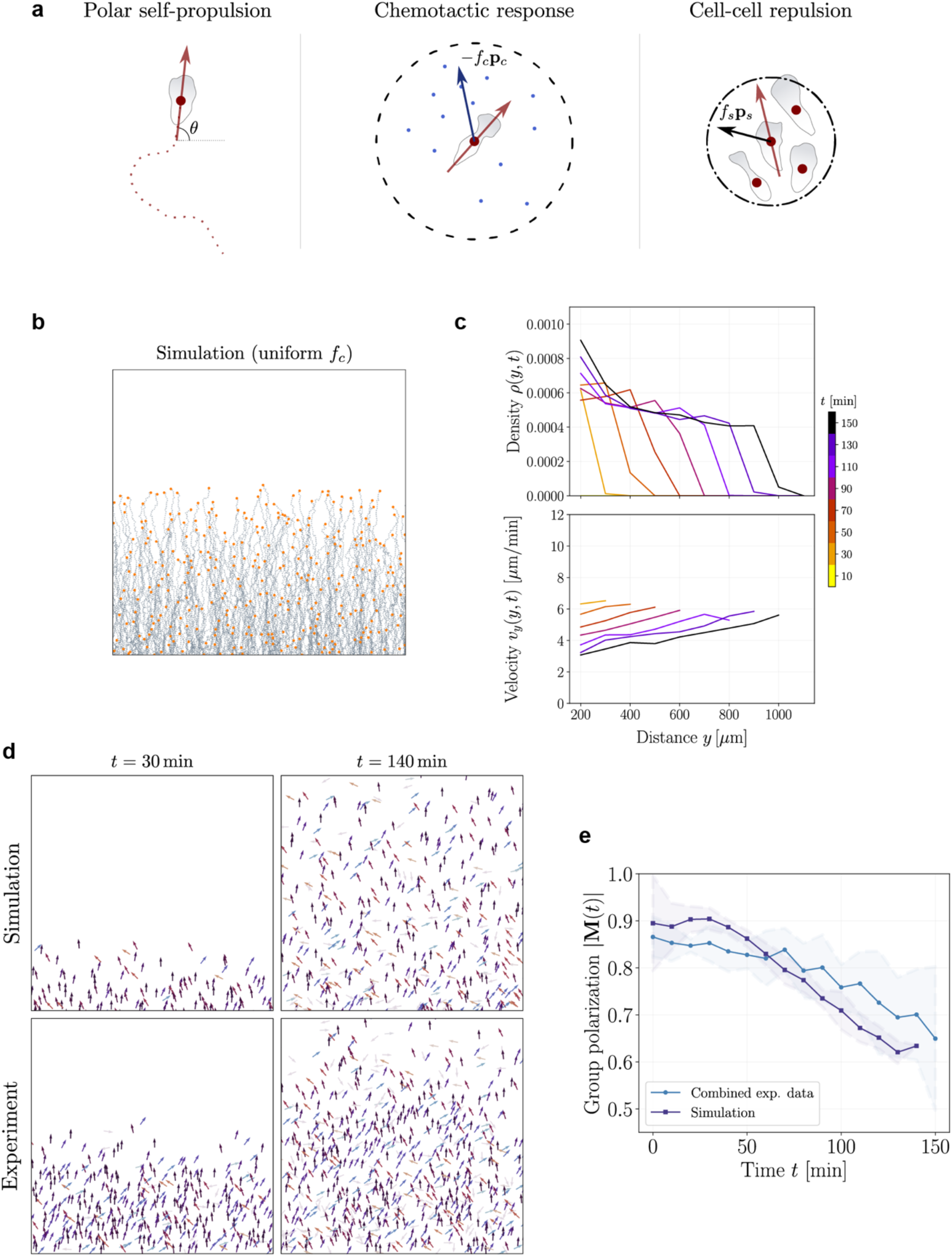
**a)** Schematics for the model assumptions used in the particle-based simulation setup. (Left) Each cell is modeled as a self-propelled polar particle with polarity (red arrow) defined by its local angle *θ* with respect to the horizontal axis. (Middle) In the presence of chemokine molecules (blue disks) within a neighborhood of radius *R_c_* (dashed circular line) surrounding the center-of-mass of the cell, a net displacement “force” (blue arrow) controlled by the chemotactic strength *f_c_* acts to reorient and displace the cell towards larger local chemokine concentrations. Upon displacement, the cell “degrades” a given percentage of sensed chemokines from this neighborhood. (Right) Each cell additionally experiences a repulsive force (black arrow) from neighboring cells in a radius *R_s_* (dash-dotted circular line), controlled by interaction strength *f_s_*, which leads to a reorientation and displacement of the cell away from its neighbors. **b)** Exemplary cell trajectories obtained from simulations with a fixed value of *f_c_* uniformly assigned to all available cells. Although the spatio-temporal dynamics are qualitatively similar to the data, the trajectories exhibit a sharper density front than those of the experimental density profiles and with simulated densities observed in the case of varying *f_c_*. **c)** Time evolution of cell densities (top) and velocities (bottom) for uniform *f_c_* showing that densities propagate as a wave while reflecting the sharp density front, and velocities decay over time as in Fig.3b but do not capture the large velocity components for the outlier cells leading at the front. **d)** Representation of instantaneous cell polarities (arrows) at different time points from simulations (top) and experiments (bottom). Color-code on the arrows indicate the corresponding local angle values with black denoting *θ*=+90°. At large times (right panels) as the population has extensively migrated, we observe larger fraction of cells with polarities deviating from the vertical orientation pointing away from the source. **e)** Group polarization decays monotonically over time both in simulations and experimental data. Averages are taken from n=4 independent experiments and n=50 simulations where the shaded regions denote means+-SDs.

## Extended data Movie legends

**Movie 1.** Matured DCs migrating towards CCL19 source in 3D collagen (25min/s, rep of 6 movies from 3 ind exp).

**Movie 2.** dPLB cells migrating towards 25nM fMLP in 3D collagen (rep of 5 ind exp).

**Movie 3.** DCs migrating under agarose with gradient (left) or uniform (right) CCL19. Shown is nuclear stain in cyan (25min/s, rep of 5 movies from 3 ind exp).

**Movie 4.** WT (left) and CCR7-KO (right) DCs migrating in PDMS device with uniform CCL19. Shown is a nuclear stain in cyan together with brightfield (10min/s, rep of 9 (WT) movies from 3 ind exp, 3 (CCR7-KO) from 2 ind exp).

**Movie 5.** Lifeact-GFP expressing DCs (cyan) migrating together with TAMRA stained WT (left) or CCR7-KO (right) T cells (magenta) under agarose with uniform CCL19 (12min/s, representatives of 6,5 from 3 ind exp).

