## Supplementary Theory Note for "Sinking the way: a dual role for CCR7 in collective leukocyte migration"

### Supplementary Theory Note - Sinking the way: Dual role of CCR7 in collective leukocyte migration

In this Supplementary Theory Note, we provide a detailed description of the particle-based simulations for cells migrating towards chemokine gradients, outline key assumptions and parameters that are used in the model, as well as details on comparison between simulations and data across different experimental setups.

#### S1 Description of particle-based simulations

As a simple framework to model the collective chemotactic migration of dendritic cells (DCs), we employ an approach based on “dry” active matter [1]. Each cell (labelled $j$) can be modelled as a particle with a given polarity (orientation) $\mathbf{e}_j \equiv (\cos(\theta_j), \sin(\theta_j))$, where $\theta_j$ is the *local angle* between the polarity vector and the (horizontal) $x$-axis. This polarity vector will then be influenced by the concentration gradients of chemokines surrounding the cell, as well as other neighboring cells and stochastic noise. In the following we outline the contribution of each of these factors.

##### S1.1 Movement of cells in the absence of chemokines

In the absence of any chemokines, each cell performs a persistent random walk of step sizes $\ell$ with a noise that induces a small rotational diffusion on the cell polarity vector $\mathbf{e}_j$. Because of the experimental observation that in the absence of chemokines, DCs only exhibit small random displacements, we use a small step size of $\ell = 0.1$ in simulation units (corresponding to a step size of $\sim 1\,\mu$m in rescaled units) for simulated cells which leads to a slow, unbiased propagation of DCs without chemokines (see Fig.3a in the main text). Each step is taken at an elementary time interval of $\tau = 1$ which correpsonds to $\Delta t = 1\,$min in rescaled units for comparison with experimental data. We implement the rotational noise of self-propulsion by changing the local angle $\theta_j$ of the cell by a small angle $\Delta\theta_j$ drawn from a uniformly distributed value within the range $[-\pi/10, \pi/10]$, i.e. at each time step the polarity vector of the cell will be tilted by a small amount. The cell then subsequently performs a step along this new polarity vector with local angle $\theta_j + \Delta\theta_j$. In real units, this rotational noise then indicates an approximate persistent time of around $\tau_p \simeq 60\,$min -defined as the characteristic time for which the polarity "forgets" a given orientation. We note that, the choices of $\ell$ and $\Delta\theta$ have a minimal influence on the simulation dynamics due to the comparatively much larger contributions arising from chemical sensing and cell-cell interactions on cell angles and coordinates.

#### S1.2 Local sensing and degradation of chemokines

We now proceed to model cells migrating in the presence local chemokine gradients. To describe their motion towards increasing chemokine concentrations, we first define a radius $R_c$ for the cells to scan for existing chemoattractants in their neighborhood. The cell displacement as a function of surrounding chemokines can then be implemented by defining a chemotactic force that will act to displace the center-of-mass of each cell. Assuming that the cell at position $\mathbf{r}$ senses $n_c$ chemokines within the radius $R_c$, we can construct a vector via the sum of distance vectors between the center-of-mass of the cell and $n_c$ chemokine molecules as:

$$\mathbf{p}_c \equiv \frac{\sum_k^{n_c}(\mathbf{r}-\mathbf{r}_k^c)}{|\sum_k^{n_c}\mathbf{r}-\mathbf{r}_k^c|} \quad \text{with} \quad |\mathbf{r}-\mathbf{r}_k^c| < R_c\,, \tag{S1}$$

where $\mathbf{r}_k^c$ is the position of the $k$-th chemokine within $R_c$. This unit vector $\mathbf{p}_c$ is then directed towards the region of *smallest* concentration of chemokines. Rescaling it by a factor $f_c$, which controls the strength/magnitude of this chemotactic sensing, we can define a chemotactic "force" $\mathbf{F}_c \equiv -f_c\mathbf{p}_c$ that points at the *largest* chemokine concentration for positive $f_c > 0$, see Ext. Data Fig.3a. Due to the chemotactic force $\mathbf{F}_c$, the cell will then be displaced to its new position given by $\mathbf{r}' = \mathbf{r} + \mathbf{F}_c$. Note that, this setup in general allows each cell with label $j$ to have a different chemotactic sensitivity $f_c^j$ and it is thus possible to explore cell populations with heterogeneous chemotactic responses. Experimental trajectories indeed indicated such a variation in individual cells' responses as inferred from their dynamic trajectories (exhibiting a mixture of fast and slow cells). We therefore both explored the settings of cell populations with homogeneous or heterogeneous chemotactic sensitivities, and found that the latter case showed a better agreement with experimental data. Indeed, we did not observe any "outlier" cells that could migrate ahead of the density front when we used a uniform/fixed value of $f_c$ for all cells, see Ext. Data Fig.3b,c for the cell trajectories, as well as density and velocity profiles obtained for uniform $f_c$.

Note that, the normalization factor (denominator) in Eq. (S2) enforces that the displacements are independent of the *absolute* concentration of chemokines: Large or small numbers of chemical molecules in the local neighborhood can lead to the same unit vector $\mathbf{p}_c$ if they exhibit the same relative concentration differences. An alternative strategy would be to use the non-normalized vector $\tilde{\mathbf{p}}_c \equiv \sum_k^{n_c}(\mathbf{r}-\mathbf{r}_k^c)$ if we wanted the chemotactic force to be dependent on the *absolute* number of locally sensed chemokines. However, at extremely saturated conditions (very large numbers of chemokines), this choice will then lead to vectors $\tilde{\mathbf{p}}_c$ that are orders of magnitude larger than the cell radius $R$, i.e. $|\tilde{\mathbf{p}}_c| \gg R$, and therefore to abrupt non-local jumps of the cell. One way to circumvent this problem is by setting a cutoff length $R_{\max}$ for these jumps that will then effectively set an upper bound for cell displacements.

To implement the degradation of chemokines surrounding the cells, we can now select a certain subset from the $n_c$ locally sensed chemokines and remove them from the simulation. For this, the two simplest choices are either removing (i) a constant number $n_d \leq n_c$, or (ii) a constant percentage $\phi_d$ of chemokines. The former method will lead to minor changes in the local chemokine gradient in regions of high absolute chemokine concentrations, because of the typically small number $n_d$ of removed particles compared to a large $n_c$ in saturated conditions. The second method, on the other hand, induces the same relative changes in gradient regardless of the absolute number of chemokines, but it might not be a realistic representation of the chemokine uptake by the cells because there is presumably an upper bound on the number of chemokine internalization. We will therefore allow cells to degrade a constant percentage of locally sensed chemoattractants, but with an upper threshold to set a maximal number $c_{max}$ of molecules that can be internalized by each cell.

##### S1.3 Chemokine diffusion and turnover

To describe the diffusion of the chemokine CCL19, we turn to a simple 2D random walk, where the coordinates of each chemical molecule is changed by $x \to x + \Delta x$ and $y \to y + \Delta y$ at each time step $\tau$. The displacements $\Delta x$ and $\Delta y$ are sampled from a uniform distribution with range $[-\ell_c, \ell_c]$, where $\ell_c$ is the maximal jump size of a chemokine molecule per time step. By varying the maximal jump size $\ell_c$ we can thus explore different regimes for the effective diffusion coefficient of the chemokine. We obtain an estimate for the maximal jumps from the diffusion coefficient of CCL19 using the relation $\tilde{D}_c \simeq \frac{1}{2\tau}\ell_c^2$, where $\tilde{D}_c$ is estimated using intensity data from FRAP experiments, see supplementary section S3 below. Finally, because in the under-agarose assays the chemokine molecules are able to diffuse in and out of the confined 2D space where cells migrate, we implemented an effective turnover at each time step by removing a certain percentage $\phi_c$ of existing chemokines from the simulation box while adding a fixed number $c_+$ of new chemokines into the simulation frame at random coordinates (determined from a uniform distribution in the range of box dimensions).

##### S1.4 Repulsion force between cells

Because the cells in the experiment presumably exhibit some type of interaction such as adhesion or volume exclusion, we wished to implement such an interaction mechanism. One simple choice would be that each cell scans a certain radius $R_s$ around its center-of-mass and will be pushed away from other neighboring cells within this region. Similar to the implementation of the chemotactic force, we could then define a vector

$$\mathbf{p}_s \equiv \frac{\sum_k^n (\mathbf{r} - \mathbf{r}_k)}{|\sum_k^n \mathbf{r} - \mathbf{r}_k|} \quad \text{with} \quad |\mathbf{r} - \mathbf{r}_k| < R_s\,, \tag{S2}$$

from the distances between the center-of-mass of each cell and of its neighbors, and rescale it by a parameter $f_s$ to obtain a net repulsion force $\mathbf{F}_s = -f_s \mathbf{p}_s$ that points *away* from the neighboring cells for $f_s < 0$. When this force acts on the cell, its new position will then become $\mathbf{r} - f_s \mathbf{p}_s$, see Ext. Data Fig.3a for an illustration of the repulsion force.

Note that, choosing a single radius for the cell size, chemokine sensing and repulsion area, i.e. $R = R_c = R_s$, corresponds to the simplest representation of a spherical cell that locally senses the chemokines "on its membrane", as well as repulsively interacts with neighboring cells upon contact. However, the experimental trajectories as well as cell shape quantifications implied that sensing and cell-cell interactions in fact were manifested at rather different length scales (i.e. indicating $R_s \neq R_c$), which we then estimated separately, see section S3 below for details. This set of rules then provides a minimal model to effectively describe individual cell motility and cell-cell interactions, and thereby to test the influence of the chemokine concentration without addressing more involved mechanisms classified under contact regulation of locomotion [2] or cell-intrinsic mechanisms such as velocity regulation [3]. Note that, this is a suitable framework if we can assume that the chemotactic as well as self-propulsion displacements of cells moving on the substrate dominate over hydrodynamic interactions mediated via diffusing molecules in the solution.

#### S2 Initialization of simulation and comparison with experiments

To initialize the simulation, we first define a simulation box with dimensions $L_x \equiv x_{max} - x_{min}$ and $L_y \equiv y_{max} - y_{min}$, and set an initial number $c_0$ of chemokine molecules with randomly distributed coordinates (from uniform distributions along the box dimensions) in the frame. Cells enter the simulation frame at constant rate during the simulation. We use periodic boundary conditions along the horizontal borders at $x_{min}$ and $x_{max}$. At the vertical borders we define reflecting boundary conditions for the chemokine molecules, but eliminate cell coordinates that leave the frame from $y_{min}$ or $y_{max}$. Due to the constant influx of cells that enter the frame during the simulation, however, the latter choice does not influence the final statistics on the trajectories. At time $t' = 0$, a fixed number $N_0$ of cells are provided at $y_{min}$ with random $x$ coordinates and polarities $\theta_j$ drawn from a uniform distribution in $[0, \pi]$. The initial number of cells are then added every $2\tau$ to represent the constant cellular influx as in the experiments. The choice of the initial number $N_0$ of cells could then be used as a parameter to explore different density regimes as explored in Fig.3c in the main text. At every time step, the positions of cells and chemokines, as well as the values of cell polarities are then updated using the rules for the sensing, degradation, and diffusion of chemokines, as well as persistent random walk of cells and cell-cell repulsions, as described in section S1 above. Note that, because the simulation boundary at $y_{min}$ defines a sharp chemokine gradient (i.e. no chemokines at $y < y_{min}$), we concentrate on cell trajectories sufficiently far from the entry region. The simulation area used for analysis then defines a square region of size $[L_x = 100, L'_y = 100]$. In rescaled units, this choice then corresponds to a region of size $1000 \times 1000 \mu\text{m}^2$ to analyze the cell trajectory data from experiments, i.e. all simulation lengths can be rescaled by a factor of 10 to obtain the corresponding experimental comparisons in $\mu$m.

To compare metrics such as density, velocity and polarity of cells, it is instructive to consider vertical segments of size $\Delta y$ spanning the considered simulation area and quantify how these metrics change over time in given $y$-segments. This projection on the vertical axis is a rather natural choice to describe the dynamics away from the cell source. At every time point $t'$, we can then determine the density of cells in a given vertical region of size $\Delta y$ by $\rho(\Delta y, t') \equiv N_{\Delta y, t'}/L_x \Delta y$, where $N_{\Delta y, t'}$ denotes the number of cells with $y$-coordinates in the considered vertical region. Similarly, we define the $y$-component of the velocity of each cell $j$ by its vertical displacements over time as $v_y^j(\Delta y, t') \equiv \delta y_j / \Delta t$, where $\Delta t = t' - t''$ is a small time interval and $\delta y_j \equiv y_j(t') - y_j(t'')$ is the vertical displacement of the $j$-th cell between $t''$ and $t'$. Note that, to obtain well-defined values for the cell velocities we consider time intervals of $\Delta t = 5$min for the experimental data (corresponding to $\Delta t = 5\tau$ in simulation units). The projected cell velocity within a vertical region $\Delta y$ at time $t'$ is then calculated by $v_y(\Delta y, t') \equiv \langle v_y^j(\Delta y, t') \rangle_{N \Delta y, t'}$ where the average is taken over all cells in the region.

#### S3 Estimation of parameter values for the simulations

Here we provide details on the parameter estimates used to tune the simulation setup to allow for a comparison with experimental cell trajectories. The key parameters that predominantly influence the emergent collective migration of cells include:

- diffusion coefficient of the chemokine $D_c$,
- internalization rate determined by the percentage $\phi_d$ and maximal numbers $c_{max}$ of chemokines

surrounding a cell,

- radii for chemotactic sensing $R_c$ and cell-cell interactions $R_s$, and
- cell density controlled by the number of cells $N_0$ added.

Because in the simulations the chemotactic coupling strength $f_c$ does not influence the polarity but only the magnitude of cell displacements (i.e. step size), we first fixed this parameter such that the overall duration for the DC front to migrate across the spatial region of interest in the experiments corresponded to the same maximal time $t_{max}$ in simulation units (effectively setting the time scale of the invasion). To account for the variability in chemotactic responses of each cell, as alluded above in section S1, we then assigned each cell a value $f_c^j$ taken from a normal distribution around the fixed value of $f_c = 0.6$ with standard deviation $\sigma_{f_c} = 0.2$. For the cell-cell repulsion strength $f_s$, we used an intermediate value that precluded cells to have overlapping coordinates when faced with the same chemokine gradient, but was still weaker than hardcore repulsions, as experimental trajectories indicated that cells in crowded regions could frequently come closer with a smaller separation than their typical radius.

To set the radius of repulsion $R_s$ we therefore looked at dense $y$-segments and from the estimated number of cells $N_y$ occupying such a region we could infer an effective repulsion radius by $R_s = L_x/N_y \simeq 20\,\mu$m. Similarly, reasoning that the chemotactic sensing radius $R_c$ can be inferred by averaging over maximal elongation of cells (which is the maximal distance at which a given cell can sense the gradient at a given time point), we obtained an estimate of $R_c \simeq 50\,\mu$m, corresponding to a maximal cell length of $100\,\mu$m, see Ext. Data Fig.1e. We then obtained a cutoff length for the chemotactic displacements to avoid observing jumps of arbitrary sizes, which we estimated from the experimental step size distributions to be around $R_{max} \simeq 100\,\mu$m. To determine the cell densities, we looked at the influx of cells from the experiments and obtained a rate of $\sim 10$ new cells entering the frame of interest every 2 min, and used this number as an estimate for $N_0$.

To determine the diffusion coefficient of CCL19, we used the data from FRAP experiments with a bleached area of radius $r \simeq 19\,\mu$m. From the intensity half-life of $t_{1/2} \simeq 950\,$ms, we obtained an estimate of $\tilde{D_c} \simeq 86\,\mu\text{m}^2/s$ [4], a typical value for the diffusion of biological proteins. In simulation units, this corresponds then to a random walk with jump sizes $\ell_c = \sqrt{2\tilde{D_c}\tau} \simeq 100\,\mu$m for time steps of $\tau = 1\,$min. An alternative value for CCL19 was given in the literature to be around $D_c \simeq 140\,\mu\text{m}^2/s$ [5] indicating jump sizes of $\ell_c \simeq 130\,\mu$m, which does not markedly change our results. Indeed, to understand the sensitivity of model predictions to chemokine diffusion, we varied the maximal attractant jump sizes in the simulations systematically between $10 - 200\,\mu$m in rescaled units to explore its influence on the collective dynamics. We found that values around $\ell_c \simeq 100\,\mu$m were a sufficiently good estimate to reproduce the experimental phenomenology.

Finally, we wished to implement the internalization, as well as the turnover of the chemokines to account for the 3D diffusion of CCL19 in and out of the quasi-2D region in the under-agarose assays. To reproduce the turnover, we deleted a fraction of $\phi_c \simeq 0.03$ of existing chemokines, and added a fixed number of chemokines at every time step $\tau$, given by the fraction $c_+ \simeq 0.07$ of the initial number $c_0 = 10^5$ of chemokines that were uniformly distributed in the simulation box. To set the internalization dynamics of the chemokines, we modelled each cell to degrade a fraction of $\phi_d = 0.05$ of chemokines that they sensed within the radius $R_c$ with a cutoff number of $c_{max} = 10$.

| | | | |
|---|---|---|---|
| Chemokine diffusion coefficient $D$ [$\mu$m$^2/s$] | 86 | Chemokine degradation fraction $\phi_d$ | 0.05 |
| Chemotactic sensing radius $R_c$ [$\mu$m] | 50 | Chemotactic strength $f_c$ | 0.6, $0.5^\dagger$ (DC), $1^\dagger$ (T cell) |
| Cell-cell repulsion radius $R_s$ [$\mu$m] | 20 | Repulsion strength $f_s$ | $-0.1$ |
| Chemokine turnover fraction $\phi_c$ | 0.03 | Chemokine influx fraction $c_+$ | 0.07 |
| Cell influx rate $N_0$ [$/2\tau$] | 10 | Cell step size $\ell$ [$\mu$m] | $10, 50^\dagger$ (T cell) |

Table 1: Parameter values used in the simulations. Symbols denoted by the dagger symbol correspond to the simulation setup for the mixed population of DCs and T cells. The length units are given in rescaled units for comparison with the experiments, and the elementary time step of the simulation corresponds to the rescaled time $\tau = 1\,\text{min}$.

Overall, Table 1 summarizes the list of key parameters used in the simulations.

### S4 Angle distributions and group polarization of the cell population

As a simple metric to determine the overall bias in cell migration profiles, we looked at distributions of local angles $\theta$ obtained from cell trajectories, as shown in Fig.3d in the main text. For the simulation data, the local angles at every time step $\tau$ were recorded for each cell during the simulation run. To determine the polarities form the experimental data, we calculated the cell displacements between each time point $t'$ and $t' - \Delta t$ (with $\Delta t = 1\,\text{min}$), which defined a unit polarity vector with local angle $\theta$ w.r.t. the horizontal axis. We then sampled the obtained local angle values for all time points over the entire cell population in the boxed region of interest.

Finally, we sought to rationalize and quantify the collective alignment of cells in response to the self-generated chemokine gradients, and turned to a metric typically used to measure the global order of polar self-propelled particles such as Vicsek-type flocks [6] or schools of fish [7] denoted as group polarization. We calculate the group polarization vector of cells at time $t$ by

$$\mathbf{M}(t) \equiv \frac{1}{N}\sum_{j=1}^{N} \frac{\mathbf{v}_j(t)}{|\mathbf{v}_j(t)|}\,, \tag{S3}$$

where $\mathbf{v}_j(t)$ is the velocity vector of the $j$-th cell and $N$ is the total number of cells in the region of interest. The normalized velocity $\frac{\mathbf{v}_j(t)}{|\mathbf{v}_j(t)|}$ thus corresponds to the polarity vector $\mathbf{e}_j$ of each cell. For a collection of cells, the magnitude $|\mathbf{M}(t)|$ then defines an order parameter, which becomes close to 0 if cells do not exhibit a preferred global orientation while increasing the collective alignment (all cells moving in the same direction) leads to a value approaching 1. To quantify the collective polarity arising from self-generated gradients we calculated the order parameter $|\mathbf{M}(t)|$ at different time points as cell populations migrated away from the entry zone. Both in simulations and experimental data, we found that the group polarization decays monotonically as a function of time, with similar values in simulations and experiments across all time points, see Ext. Data Fig.3e. As early time points mainly represent the propagating front of cells, the decrease in the order parameter indicated that cells in the bulk region (behind the front) showed larger fluctuations in their polarity. Note that, this decrease of global orientational bias in the bulk is not linked to the decrease in the cell velocities as shown in Fig.3b of the main text: The metric defined in Eq.(S3) is determined by the *instantaneous polarity* of cells, and thus provides an independent measure for the impaired bias in the population bulk.

### S5 Simulation of DCs migrating in the microfluidic maze

Here we briefly describe the simulation setup that reproduces the migration of DCs in a minimal microfluidic "maze". We first constructed a geometric replica of the maze setup in the simulation box with rescaled units. In the experimental setup, cells enter the maze at the bottom from a large chamber, and the "open end" of the maze connects to a large reservoir of CCL19. To mimic this large chemokine reservoir, we extended the "open" channel of the maze to end at a large $y$ value (approx. $10\times$ the length of the "closed" channel). As an initial condition, the maze was filled uniformly with a fixed number of chemokine molecules, which could not escape the maze during the simulation due to reflecting boundary conditions at the walls of each channel. Furthermore, unlike the under-agarose setup, we omitted the turnover/decay of chemokine molecules because in this experiment a 3D diffusion of chemokines would not be possible, i.e. chemokines are now only degraded via internalization by the cells and no chemokines are added into the maze during the course of the simulation. The interaction of cells with the channel walls could be described by a repulsive force as in the case of cell-cell repulsion (while for the rare cases where some cells crossed over the maze walls despite the repulsion, we eliminated these cells from the simulation). We then initialized the cells to enter the maze from the bottom end and applied the same rules as used to simulate the under agarose assays to update cell & chemokine positions and cell polarities. We found that cells migrated preferentially towards the open end when the diffusion coefficient of the chemokine $D_c$ was sufficiently large. In fact, for small $D_c$ cells showed no preference between the open and closed channels (%50 probability to enter both), whereas the percentage of cells entering the open-end channel increased monotonically with $D_c$, with biologically relevant values of $D_c \simeq 100 - 200\,\mu\text{m}^2/s$ leading to probabilities of $\%60 - 80$ to enter the open-end channel, see Ext. Data Fig.2d. This influence of the attractant diffusibility on the cell behavior was indeed also previously found in simulations of cells in complex mazes consisting of several dead-end channels [8].

### S6 Simulation of DCs mixed with T cells

To model the experiments with two populations of dendritic and T cells co-migrating in the under agarose assay, we modified the simulation setup by labelling a small fraction of cells to represent T cells, which had a different chemotactic sensitivity than DCs and could not internalize/degrade chemokine molecules. We chose a stronger chemotactic strength $f_c^T$ for T cells (here using fixed values $f_c^T = 1$ for T cells and $f_c = 0.5$ for DCs) to reproduce the overall larger migration speed of T cells compared with that of DCs. Note that, because the chemotactic strength $f_c$ determines the magnitude of cell displacements due to local chemokine gradients, a large value of $f_c$ effectively corresponds to a large chemotactic coefficient as in mean-field models of chemotaxis.

In the simulations with mixed populations of DCs and T cells, we systematically observed that T cells propagated with a concentration maxima at a characteristic distance $\mathcal{L} \simeq 150\,\mu\text{m}$ ahead of the

DC density front at all times, see Ext. Data Fig.2f. Despite their larger chemotactic sensitivity (as controlled by $f_c^T$), T cells could not migrate further away because they lost their polarity due to the shallow chemokine gradients at distances larger than $\mathcal{L}$ from the DC front. The propagation speed of the T cell wave front was therefore strongly coupled to the speed of the DC front, which was in turn determined by the local gradient generation via chemokine internalization. To test whether this was indeed the underlying mechanism for the coupling of two waves, we modified the T cell properties in a control simulation such that they could not respond to surrounding chemokines (setting $f_c^T = 0$) but were able to take larger steps ($\ell = 0.5$) to migrate over long distances. We found that in this control setup T cells migrated "independently" of the DC concentration, without any coupling to the DC front and without a collective directional bias, see Fig.4e (right panel) in the main text.
